# Neuronal activity regulates Matrin 3 levels and function in a calcium-dependent manner through calpain cleavage and calmodulin binding

**DOI:** 10.1101/2022.04.11.487904

**Authors:** Ahmed M. Malik, Josephine J. Wu, Christie A. Gillies, Quinlan A. Doctrove, Xingli Li, Haoran Huang, Vikram G. Shakkottai, Sami Barmada

## Abstract

RNA-binding protein (RBP) dysfunction is a fundamental hallmark of amyotrophic lateral sclerosis (ALS) and related neuromuscular disorders. Abnormal neuronal excitability is also a conserved feature in ALS patients and disease models, yet little is known about how activity-dependent processes regulate RBP levels and functions. Mutations in the gene encoding the RBP Matrin 3 (MATR3) cause familial disease, and MATR3 pathology has also been observed in sporadic ALS, suggesting a key role for MATR3 in disease pathogenesis. Here, we show that glutamatergic activity drives MATR3 degradation in a NMDAR-, Ca^2+^-, and calpain-dependent mechanism. The most common pathogenic *MATR3* mutation renders it resistant to calpain degradation, suggesting a link between activity-dependent MATR3 regulation and disease. We also demonstrate that Ca^2+^ regulates MATR3 through a non-degradative process involving the binding of Ca^2+^/calmodulin (CaM) to MATR3 and inhibition of its RNA-binding ability. These findings indicate that neuronal activity impacts both the abundance and function of MATR3, and provide a foundation for further study of Ca^2+^-coupled regulation of RBPs implicated in ALS and related neurological diseases.

## Introduction

Amyotrophic lateral sclerosis (ALS) is a neurodegenerative disease involving the loss of upper and lower motor neurons, resulting in muscle weakness, paralysis, and ultimately death. ALS shares clinical, genetic, and pathological overlap with frontotemporal dementia (FTD), a disorder characterized clinically by altered behavior and speech^1^, and pathologically by the degeneration of neurons in frontal and temporal cortices. Accumulating evidence points to the dysfunction of RNA-binding proteins (RBPs) in the pathogenesis of both ALS and FTD, with the vast majority of patients with ALS and approximately half of those with FTD displaying aggregation and/or mislocalization of the RBP TDP43 in affected tissues on autopsy^2,3^. Moreover, while most cases of ALS and FTD are sporadic, rare mutations including in genes encoding RBPs can cause inherited disease^4–7^.

To date, several disease-linked RBPs have been identified, including but not limited to MATR3, TDP43, FUS, and hnRNPA2/B1. Mutations in the genes encoding these proteins can not only cause ALS and FTD, but also primary myopathy^8^,^9^. MATR3 is a nuclear protein with two zinc finger (ZF) domains and two RNA recognition motifs (RRMs) that has the capacity to bind both DNA and RNA targets^10–12^. Over a dozen point mutations in the *MATR3* gene have been linked to familial ALS, combined ALS/FTD, or hereditary distal myopathy^13–21^. Furthermore, MATR3 pathology—consisting of MATR3 mislocalization, changes in abundance, and aggregation—has been found in motor neurons of sporadic ALS patients^22,23^, suggesting a fundamental link between MATR3 dyshomeostasis and neuromuscular disease.

Abnormal neuronal activity is frequently observed in ALS and FTD patients^24–26^ as well as disease models, with both hypo- and hyperexcitability phenotypes reported depending on neuronal population studied, model system, and disease progression^27–31^. Although regulation of RBP abundance and function in response to neuronal hypo- or hyperactivity in disease is incompletely understood, several RBPs are affected by receptor activation and/or depolarization^32–35^. These observations indicate that investigations of disease-linked RBPs in the context of neuronal activity may reveal crucial disease-related mechanisms contributing to RBP pathology and perhaps neurodegeneration in ALS and FTD.

In previous studies, MATR3 was rapidly cleared from cerebellar granule neurons after treatment with *N*-methyl-D-aspartate (NMDA) in a protein kinase A (PKA)-dependent manner^36^. Separate *in vitro* studies suggested that MATR3 binds Ca^2+^-bound calmodulin (CaM)^37^, a central signal transduction factor that rapidly shuttles to the nucleus upon associating with Ca^2+^, thereby driving the expression of key plasticity-related transcriptional programs^38–42^. Even so, whether or how CaM impacts MATR3 function and levels downstream of NMDA receptor activation remains unknown.

Here, we show that neuronal MATR3 is degraded after glutamatergic stimulation through an NMDAR-, Ca^2+^-, and calpain-dependent pathway. Prior to its degradation, however, MATR3’s interaction with Ca^2+^/CaM also inhibits its ability to bind RNAs, suggesting that Ca^2+^/CaM may exquisitely tune the function of MATR3 and related RBPs through two complementary mechanisms that operate at distinct time scales.

## Results

### Glutamatergic stimulation reduces MATR3 in vitro and ex vivo

We first sought to confirm the effect of NMDAergic stimulation on MATR3 levels. Treatment of mature (DIV14-16) cortical neuron cultures with NMDA resulted in a rapid, time-dependent decrease in MATR3 protein abundance (Fig.1A-B). Notably, and in contrast to previous work in cerebellar neurons that indicated MATR3 clearance is PKA-mediated, we observed only a trend toward blunting of NMDA-dependent MATR3 decrease by the PKA inhibitor H89 (Supplemental Fig. 1A-B). Using a PKA substrate antibody, we also noted an accumulation of phosphorylated MATR3 upon NMDA treatment, but MATR3 phosphorylation was not blocked by H89 (Supplemental Fig. 1CD), suggesting that a kinase other than PKA phosphorylates MATR3 at a residue recognized by this antibody. These observations argue against PKA-mediated clearance of MATR3 in cortical neurons upon NMDA stimulation, and prompted us to examine the mechanism of MATR3 degradation in more detail.

**Figure 1.**
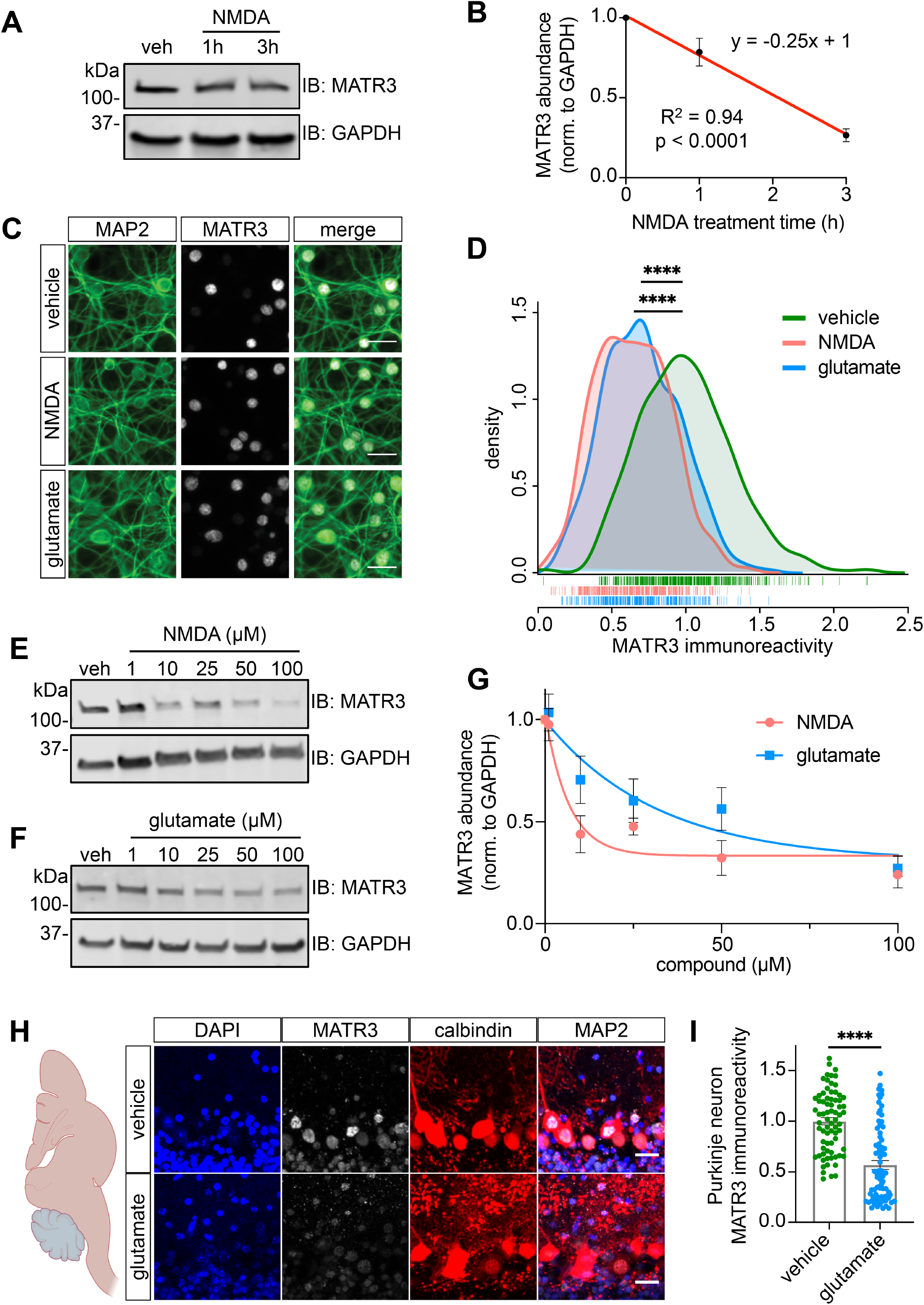
Glutamatergic stimulation triggers rapid MATR3 reduction *in vitro* and *ex vivo*. (**A,B**) Treatment of mature DIV14-16 cortical neuron cultures with 100μM NMDA results in rapid, time-dependent MATR3 decrease. (n=3; p-value determined by sum of squares F-test). (**C,D**) This reduction is recapitulated with immunostaining but is not accompanied by MATR3 redistribution within neurons (vehicle, n=547; NMDA, n=419, ****p<0.0001; glutamate, n=337, ****p<0.0001; one-way ANOVA with Tukey’s post-hoc test). (**E-G**). Both NMDA and the endogenous agonist glutamate elicit dose-dependent MATR3 clearance (vehicle, n=3 per dose, y=0.69^-0.15x^ + 0.33; glutamate, n=3 per dose, y=0.69^-0.031x^ + 0.31; nonlinear one phase decay). (**H,I**) Calbindin-positive Purkinje neurons in cerebellar slice cultures displayed markedly reduced MATR3 staining intensity upon glutamate stimulation (vehicle, n=73; NMDA, n=76, ****p< 0.0001; two-tailed t-test). Scale bars in (**C**) and (**H**), 20μm.

We first asked whether glutamate, the endogenous agonist for NMDARs as well as other glutamatergic receptors, likewise results in MATR3 decrease. Treatment with equivalent doses of NMDA or glutamate for 3h resulted in comparable reductions of MATR3 immunoreactivity in MAP2-positive neurons without MATR3 aggregation or cytoplasmic mislocalization, indicating that MATR3 clearance occurs primarily in the nuclear compartment (Fig. 1C-D). Dose-response experiments with NMDA and glutamate likewise confirmed that both compounds were able to effectively decrease MATR3 levels (Fig. 1E-G).

### Glutamatergic stimulation regulates MATR3 levels ex vivo

We questioned whether this phenomenon, which we originally identified in primary cultures of mixed cortical neurons, also regulates MATR3 levels in other settings. In particular, we focused on *ex vivo* preparations of Purkinje cells of the cerebellum, since these cells display a striking cell-to-cell heterogeneity in MATR3 abundance in basal conditions^43^ and susceptibility to mutant *MATR3*-mediated neurotoxicity^44^. To investigate MATR3 clearance in Purkinje cells, we generated acute cerebellar slices from wild-type C57BL/6J mice. These slices were then treated with vehicle or glutamate before fixation and immunostaining for MATR3 and the Purkinje cell marker calbindin (Fig. 1H). Similar to neurons isolated from cortex (Fig. 1C-D), Purkinje neurons treated with glutamate displayed markedly reduced MATR3 immunoreactivity compared to vehicle-treated slices (Fig. 1I). These data indicate that activity-dependent regulation of MATR3 applies not just to primary cortical neurons but also to Purkinje neurons of the cerebellum; moreover, this pathway may at least in part explain previous observations of MATR3 variance in Purkinje neurons.

### MATR3 is cleared in an NMDA- and Ca2+-dependent process

Neurons express a variety of receptors activated by glutamate, including the ionotropic kainate, AMPA, and NMDA receptors as well as a host of metabotropic receptors that do not directly pass Na^+^ or Ca^2+^ but instead trigger intracellular second messengers (Fig. 2A). Treatment with AP5, a selective antagonist of NMDARs, had no effect on MATR3 abundance on its own. Nevertheless, AP5 blocked MATR3 clearance not only in response to NMDA—as expected from coadministration of the agonist and the antagonist—but also in response to glutamate (Fig. 2B-C). These results indicate that NMDAR stimulation is both necessary and sufficient for glutamatergic MATR3 reduction in mixed primary cortical neurons.

**Figure 2.**
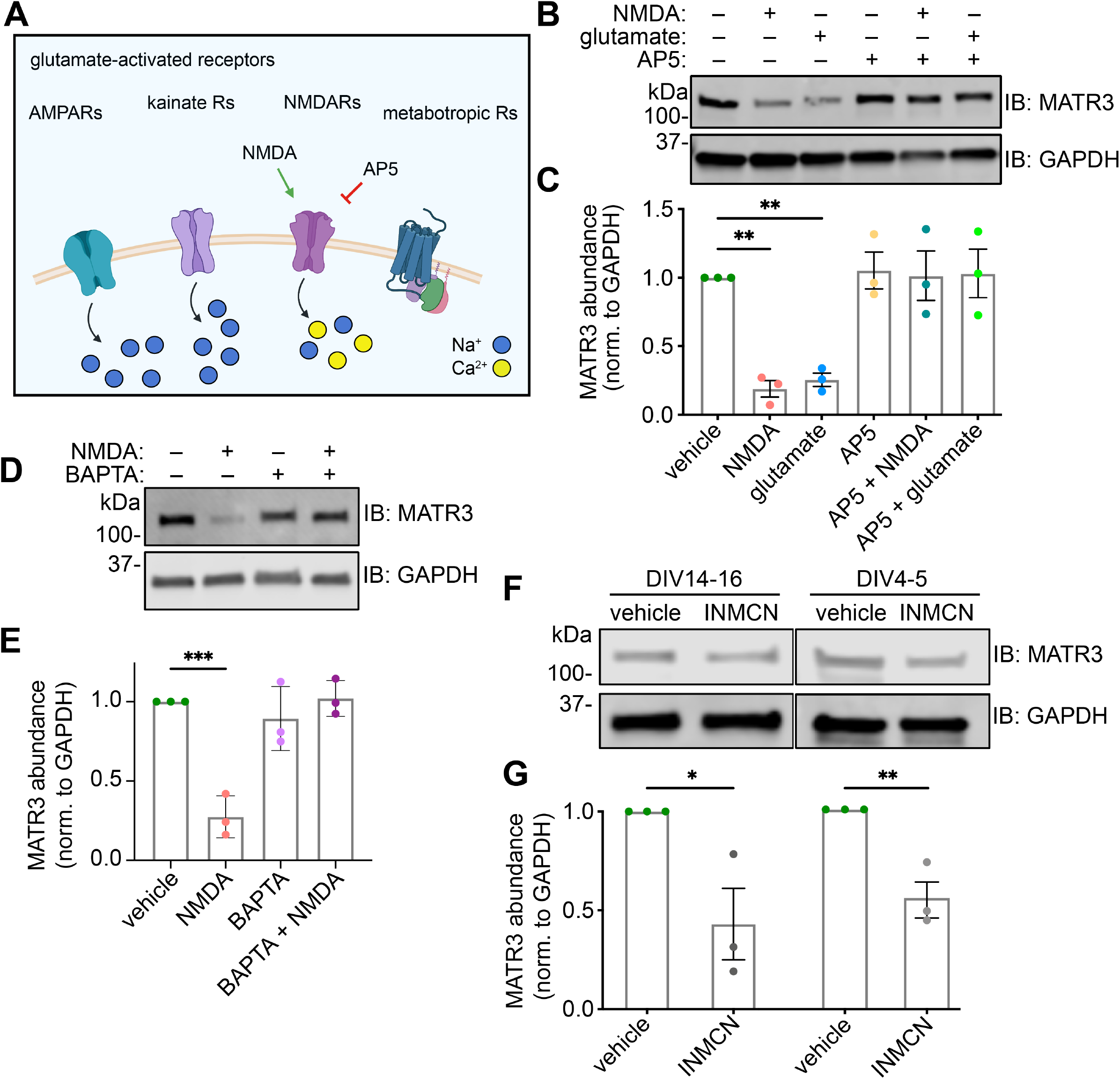
MATR3 reduction upon glutamatergic stimulation is NMDAR- and Ca^2+^-dependent. (**A**) Neurons express a variety of ionotropic and metabotropic receptors activated by glutamate, with NMDA being a selective agonist and AP5 a selective antagonist of NMDARs. Unlike other glutamate receptors, NMDARs are largely unique in passing Ca^2+^ when activated. (**B,C**) NMDAR activation is both necessary and sufficient for MATR3 clearance (n=3; **p<0.01; ns, not significant; one-way ANOVA with Tukey’s post-hoc test). (**D,E**) The Ca^2+^ chelator BAPTA blocks NMDA-mediated MATR3 reduction, indicating that Ca^2+^ influx is necessary for this process (n=3; ***p<0.001; ns, not significant; one-way ANOVA with Tukey’s post-hoc test). (**F,G**) Increasing intracellular Ca^2+^ with ionomycin (INMCN) is sufficient for MATR3 reduction in mature DIV14-16 (n=3; *p=0.034, two-tailed t-test) and immature DIV4-5 (n=3; *p=0.0080, two-tailed t-test) cultures.

Canonical NMDARs are largely unique among the main classes of glutamatergic receptors in being permeable to both Ca^2+^ or Na^+^, and so we wondered whether the dependence of MATR3 clearance on NMDARs stems from this process being dependent on Ca^2+^. To answer this, we treated neurons with the Ca^2+^ chelator BAPTA before NMDA treatment and found that this chelation blocked NMDA-induced reduction in MATR3, indicating that MATR3 clearance requires Ca^2+^ entry into the cell (Fig. 2D-E). If an increase in intracellular Ca^2+^ is the mechanism downstream of NMDAR activation that drives MATR3 reduction, we would expect non-physiological increase in intracellular Ca^2+^ independent of receptor engagement to mimic this effect. Indeed, treatment with the ionophore ionomycin resulted in MATR3 protein decrease not only in NMDAergically mature DIV14-16 neurons (Fig. 2F-G) but also in NMDAergically immature DIV4-5 cells (Fig. 2H-I), indicating that increased intracellular Ca^2+^ is both necessary and sufficient for MATR3 reduction. Notably, ionomycin treatment in HEK293T and Neuro2A cell lines did not result in MATR3 decrease, suggesting that the Ca^2+^-dependent mechanisms responsible may rely on neuron-specific or neuron-enriched factors (Supplemental Fig. 2A-B).

### NMDA-related reduction in MATR3 occurs post-transcriptionally via calpain-mediated cleavage

The reduction in MATR3 we observed upon NMDAR stimulation could be due to a decrease in the production of MATR3, degradation of existing MATR3, or both. Using RT-PCR, we observed only a weak trend towards reduced *Matr3* production upon NMDA stimulation (Fig. 3A), which—along with the rapid timeframe of reduction (Fig. 1B)—points to degradation of MATR3 rather than robust transcriptional downregulation as the primary mechanism for its reduction. In a complementary experiment, we transduced cultures with lentivirus expressing FLAG-tagged MATR3. This construct expressed from exogenous cDNA would be expected to be unaffected by transcriptional or post-transcriptional mRNA control, and thus any NMDA-induced changes in this protein’s abundance would come from post-translational degradation. Indeed, NMDA treatment resulted in the rapid reduction of this protein as determined by probing with a FLAG antibody (Fig. 3B), supporting a model in which NMDAR stimulation triggers the clearance of existing MATR3 protein.

**Figure 3.**
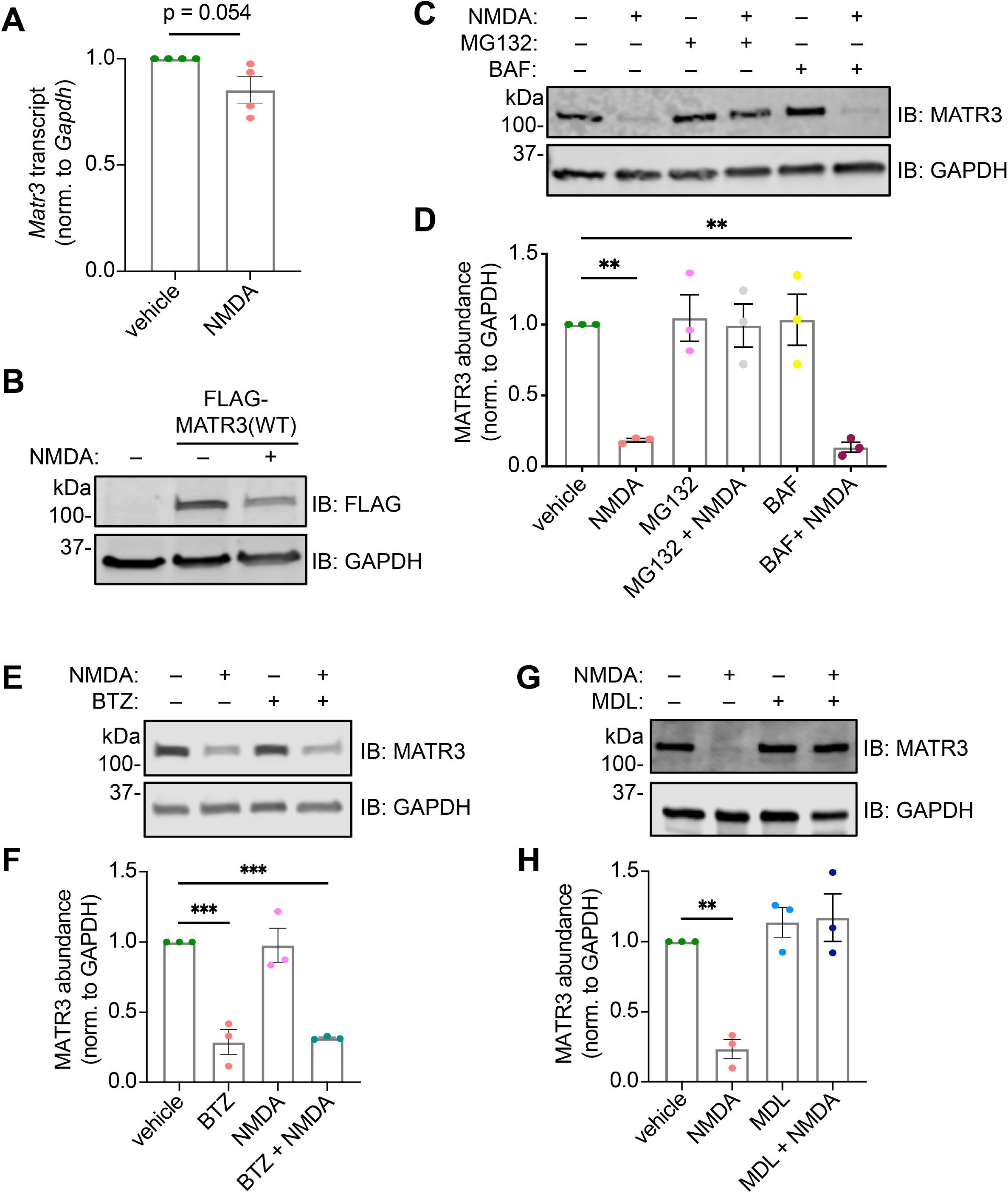
NMDA-triggered MATR3 reduction is accomplished post-translationally via calpains. (**A**) Treatment with NMDA does not significantly alter *Matr3* transcript abundance (n=4; p=0.054, two-tailed t-test). (**B**) Abundance of exogenous, lentivirally delivered FLAG-MATR3 is reduced by NMDA application. (**C**) The proteasomal and cysteine protease inhibitor MG132, but not the autophagy inhibitor bafilomycin (BAF), blocks MATR3 clearance in response to NMDA (n=3; **p=0.0041; p * 0.0025; one-way ANOVA with Tukey’s post-hoc test). (**E,F**) The selective proteasomal inhibitor bortezomib (BTZMB) fails to block NMDA-triggered MATR3 reduction (n=3; ***p<0.001, one-way ANOVA with Tukey’s post-hoc test). (**G,H**) MDL28170 (MDL), an inhibitor of calpains, effectively impairs MATR3 degradation upon NMDA treatment (n=3; **p=0.0040; one-way ANOVA with Tukey’s post-hoc test).

We then investigated the pathway responsible for NMDA-related MATR3 degradation. MATR3 turnover in response to NMDA is completely blocked by MG132—a broad inhibitor of cysteine proteases as well as threonine proteases that constitute the proteasome^45^—but not the vacuolar ATPase blocker bafilomycin^46^ (Fig. 3B-D), suggesting that MATR3 degradation requires the ubiquitin-proteasome system (UPS) or cysteine proteases. Subsequent experiments demonstrated that bortezomib, a specific inhibitor of the 26S proteasome subunit^47^, failed to prevent MATR3 degradation in response to NMDA, indicating that an MG132-sensitive protease independent of the UPS is responsible for MATR3 cleavage (Fig. 3E-F). Further supporting this conclusion, we saw no accumulation of higher molecular weight, ubiquitinated MATR3 species in neurons treated with MG132 and NMDA (Supplemental Fig. 2C). Therefore, we asked whether calpains, a group of Ca^2+^-activated, MG132-sensitive cysteine proteases, may be cleaving MATR3. Treatment with MDL28170, a specific calpain inhibitor, effectively blocked NMDA-induced MATR3 degradation (Fig. 3G-H), pointing to calpains as the effector proteases of NMDA-mediated degradation. We were unable to detect lower molecular weight MATR3 species after NMDA treatment, even in the setting of proteasomal inhibition and using antibodies targeting both N- and C-termini of MATR3, indicating that these fragments are rapidly cleared in a UPS-independent manner (Supplemental Fig. 3A-B). These results are in alignment with previous reports demonstrating the *in vitro* susceptibility of MATR3 to calpains^48^, and the apparent difficulty in detecting calpain cleavage products^49,50^

The two calpains most expressed in the nervous system are calpain 1 (CAPN1) and calpain 2 (CAPN2), with CAPN1 enriched in neurons and CAPN2 expressed predominantly by glia^51,52^. In previous studies, CAPN1/2 overexpression in HEK293T cells resulted in calpain activation even without the addition of Ca^2+^ due to low expression of the endogenous calpain inhibitor calpastatin, elevated basal Ca^2+^ concentrations, or a combination of these factors1^53,54^. Consistent with this, increasing Ca^2+^ in HEK293T cells by application of the ionophore ionomycin had little effect on MATR3 levels (Supplemental Fig. 2A). However, CAPN1 overexpression significantly reduced endogenous MATR3 levels in transfected HEK293T cells, compared to controls (Fig. 4A-B). In these experiments, CAPN1 but not CAPN2 stimulated MATR3 clearance, suggesting that CAPN1 selectively degrades MATR3.

**Figure 4.**
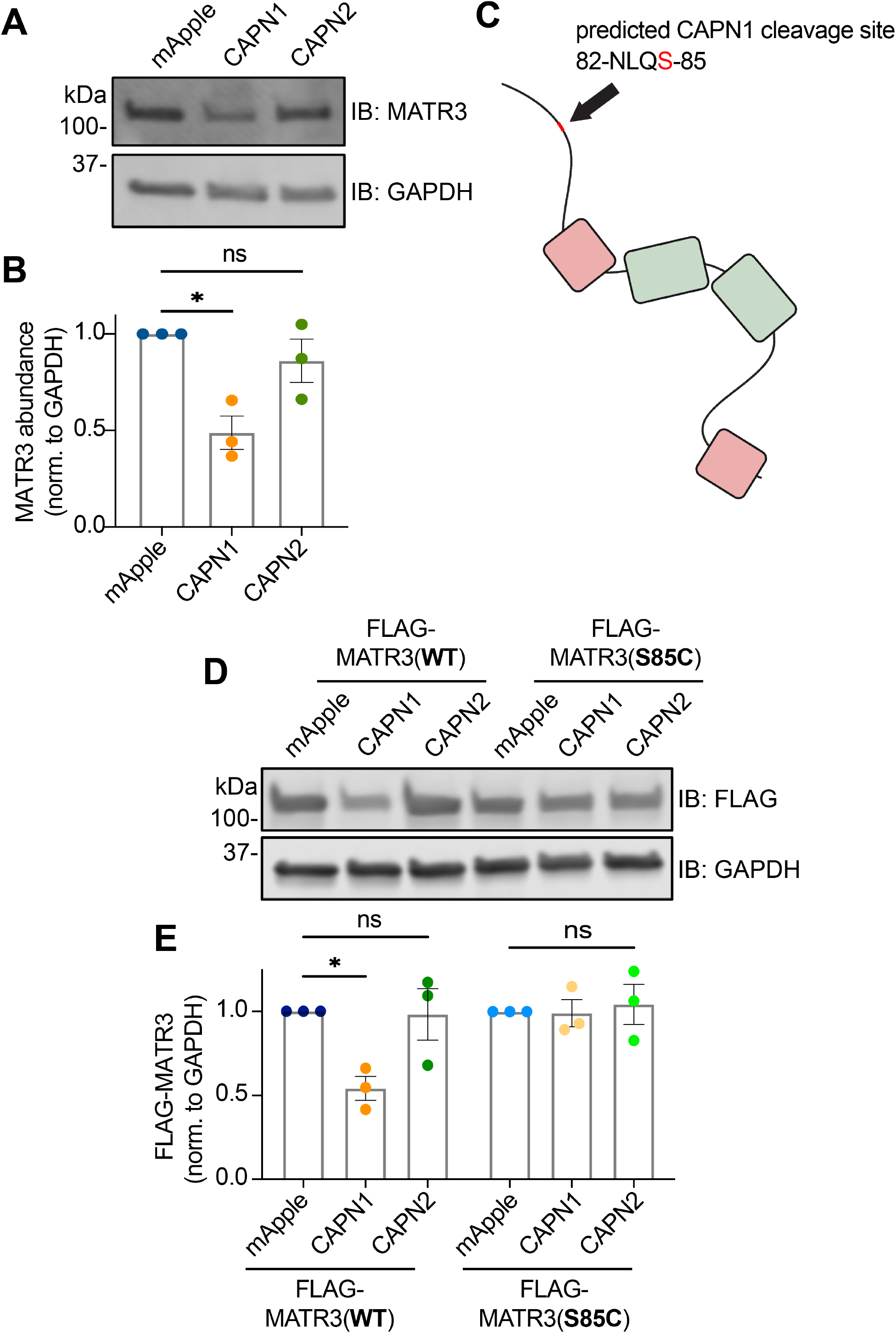
MATR3 is a substrate for CAPN1, and the pathogenic S85C mutation renders it resistant to degradation. (**A,B**) CAPN1 but not CAPN2 overexpression resulted in MATR3 degradation in HEK293T cells (n=3; *p=0.10, one-way ANOVA with Tukey’s post-hoc test). (**C**) The highest confidence CAPN1 cleavage site in MATR3 spans amino acids 82-85; notably this includes the S85 residue (red) mutated in familial disease. (**D,E**) While exogenous FLAG-MATR3(WT) is susceptible to cleavage by CAPN1 (n=3; *p=0.036; ns, not significant; one-way ANOVA with Tukey’s post-hoc test), the pathogenic S85C mutation is resistant (n=3; one-way ANOVA with Tukey’s post-hoc test).

An online calpain substrate algorithm^55^ predicted a strong candidate for a CAPN1 cleavage site at residues 82-85 of MATR3 (Fig. 4C). This stretch of amino acids includes S85, a residue altered by the most common pathogenic *MATR3* mutation (S85C). Therefore, we asked whether this mutation affects MATR3 cleavage by CAPN1. In HEK293T cells transfected with mApple, CAPN1, or CAPN2 and FLAG-tagged MATR3 variants, exogenous FLAG-MATR3(WT) was degraded by CAPN1 to a comparable degree as endogenous MATR3. In contrast, FLAG-MATR3(S85C) was completely resistant to CAPN1-mediated cleavage (Fig. 4D-F). Taken together, these data suggest that NMDAR activation leads to MATR3 degradation through a Ca^2+^- and calpain-dependent manner, a process that is impaired by the disease-linked S85C mutation.

### Ca^2+^ promotes a selective interaction between Ca^2+^/CaM and MATR3

Our results indicate that elevated Ca^2+^ over the course of hours drives MATR3 clearance. Even so, we wondered whether stimulation on shorter, more physiologic timescales could also tune MATR3 function. Beyond activation of calpains, Ca^2+^ may impact several downstream pathways in neurons, many of which are modulated via calmodulin (CaM), a ubiquitous and highly conserved factor. Upon binding Ca^2+^ ions, CaM undergoes conformational changes that greatly alter its affinity for other proteins, resulting in their activation or inhibition. Prior studies^37^ suggested that MATR3 binds selectively to Ca^2+^/CaM via a sequence overlapping the MATR3 RRM2. Using publicly available informatics algorithms^56–58^, we confirmed the presence of a predicted CaM-binding site on MATR3 within RRM2, corresponding to the MATR3 fragment used in previous studies (Fig. 5A)^37^. We also identified analogous CaM-binding sites buried within the RRMs of several RBPs linked to ALS/FTD, suggesting that CaM-RBP interactions may extend beyond MATR3.

**Figure 5.**
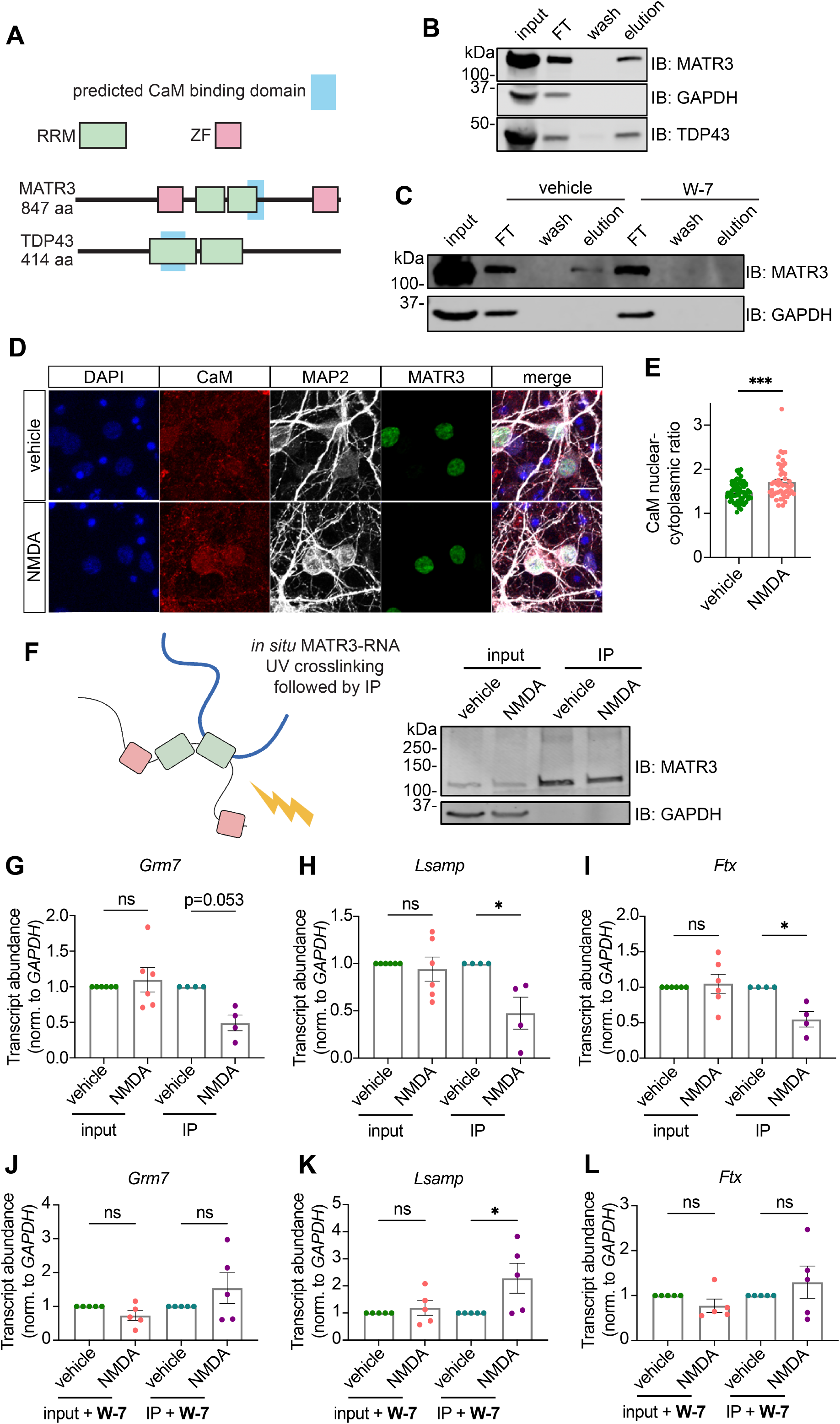
CaM binds to MATR3 in a Ca^2+^-sensitive manner and interferes with recognition of RNA by MATR3. (**A**) CaM-binding motif prediction algorithms indicate that CaM interacts with the RNA recognition motifs (RRMs) of both MATR3 and TDP43. (**B**) Ca^2+^-bound but not free apo-CaM binds to MATR3 and TDP43, validating *in silico* predictions. (**C**) The conformational CaM inhibitor W-7 blocks the association between Ca^2+^/CaM and MATR3. (**D,E**) Ca^2+/^CaM is enriched within the nucleus of neurons treated with NMDA (vehicle, n=67; NMDA, n=48; ***p<0.0001; two-tailed t-test). Scale bars in (**D**), 20μm. (**F**). UV crosslinking followed by IP (UV-CLIP) was used to analyze RNA targets bound to MATR3 *in situ*. (**G-I**) While NMDA treatment did not alter total levels of MATR3 target RNAs (n=6 for each candidate), less RNA was crosslinked to MATR3 in stimulated conditions, indicating impaired RNA binding (n=4 for each candidate; *p<0.05, two-tailed t-test). (**J-L**) Pre-treatment with CaM inhibitor W-7 prior to NMDA stimulation did not alter total levels of MATR3 target RNAs (n=5 for each candidate). Although W-7 pretreatment increased *Lsamp* RNA crosslinked to MATR3 (*p=0.045, two-tailed t-test), it did not alter the amount of *Grm7* or *Ftx* crosslinked to MATR3 (n=5 per condition; ns, not significant, two-tailed t-test).

To validate these predicted binding events, we incubated lysates from HEK293T cells with CaM-conjugated sepharose beads in the presence of Ca^2+^. Following washing, elution was accomplished by incubation with a Ca^2+^ chelator, stripping Ca^2+^ from Ca^2+^/CaM, returning CaM to its apo conformation, and selectively liberating those proteins bound to Ca^2+^/CaM. In this way, we found that both MATR3 and TDP43 bind CaM in a Ca^2+^-dependent manner, recapitulating previous findings on MATR3 while also suggesting that CaM may regulate additional disease-associated RBPs (Fig. 5B). Focusing our studies on MATR3, we sought to further confirm the selectivity of its interaction with CaM. For this, we incubated lysates with CaM-conjugated beads in the absence or presence of W-7, a small molecule inhibitor that impairs CaM’s ability to adopt its activated confirmation even in the presence of Ca^2+^. W-7 completely blocked MATR3 binding to CaM (Fig. 5C), indicating that Ca^2+^ does not promote the CaM-MATR3 interaction through any effects on MATR3 but rather through the Ca^2+^-triggered conformational changes in CaM.

### Activity rapidly impairs MATR3 binding to substrate RNAs

Given the location of the CaM-binding site within the MATR3 RRM2 domain, we wondered whether CaM binding downstream of Ca^2+^ influx could interfere with the ability of MATR3 to recognize its RNA substrates. CaM is a small protein capable of passing into the nucleus by passive diffusion and thus is found distributed diffusely in the cell. Upon neuronal depolarization, however, CaM rapidly translocates to the nucleus, where it drives the expression of activity-dependent genes^38–42^. In keeping with these results, NMDA treatment of primary neurons results in a modest increase in nuclear CaM within a matter of minutes (Fig. 5D,E). This effect, along with its adoption of the activated Ca^2+^-bound conformation, would be expected to enhance its association with resident nuclear proteins such as MATR3.

To determine if CaM binding displaces RNA from MATR3, we irradiated primary neurons after 5 min of NMDA treatment, thereby crosslinking RNA-MATR3 complexes. We then immunoprecipitated MATR3, isolating bound high-confidence neuronal RNA targets identified in previous studies^59^ and measuring their abundances by quantitative RT-PCR (UV CLIP; Fig. 5F). MATR3 pulled down significantly less crosslinked RNA in the context of NMDA stimulation, compared to vehicle control, despite no change in the overall abundance of each target (Fig. 5G-I). Furthermore, pre-treatment with the CaM inhibitor W-7 prevented the NMDA-induced drop in MATR3 binding for these targets (Fig. 5J-L), and in the case of one MATR3 substrate (*Lsamp*), W-7 pretreatment led to an increase in MATR3 binding. These results suggest that NMDA stimulation impairs RNA recognition by MATR3 through a Ca^2+/^CaM-dependent manner, affecting the function of MATR3 without directly reducing its abundance.

We also wondered whether CaM activation may have more sustained effects beyond inhibiting MATR3 RNA binding. Indeed, CaM inhibition with W-7 completely blocked MATR3 degradation in response to NMDA (Supplemental Fig. 4A-B). In light of these results and our previous data suggesting destabilization of neuronal TDP43 upon interruption of RNA binding^60^, we asked whether CaM activation might likewise promote MATR3 clearance by inhibiting its RNA binding. To answer this, we turned to optical pulse labelling (OPL), a technique in which proteins of interest are fused to the photoconvertible protein Dendra2 and expressed in primary neurons. Brief illumination with UV light triggers the irreversible conversion of Dendra2 from a green fluorescent protein to a red fluorophore. The time-dependent loss of red fluorescence for each cell can then be used to determine a half-life for Dendra2-tagged proteins. Using this method, we observed no differences in the turnover of MATR3(WT)-Dendra2 and a variant of MATR3 with its key RNA-binding domain deleted (ΔRRM2). These data suggest that, in contrast to TDP43, impaired RNA binding *per se* does not destabilize MATR3.

## Discussion

In this work, we focused upon the physiological regulation of MATR3 abundance and function in neurons. Application of glutamate resulted in the rapid clearance of MATR3 in an NMDA-dependent manner. In contrast to previous observations, however, PKA inhibition only produced a trend towards blunted MATR3 clearance upon NMDA treatment. Both Ca^2+^ and calpains were required for neuronal MATR3 reduction in response to NMDA, with CAPN1 being both necessary and sufficient for MATR3 clearance. Based upon the predicted location of the CAPN1 cleavage site, we found that the most common pathogenic MATR3 mutant, S85C, renders the protein resistant to CAPN1 cleavage (Fig. 6). Together with our prior data showing that MATR3 levels are critical determinants of neuronal survival^61^, these findings suggest that interference with physiological regulation of MATR3 function and abundance may represent a novel pathogenic mechanism underlying S85C-induced neurodegeneration and myopathy.

**Figure 6.**
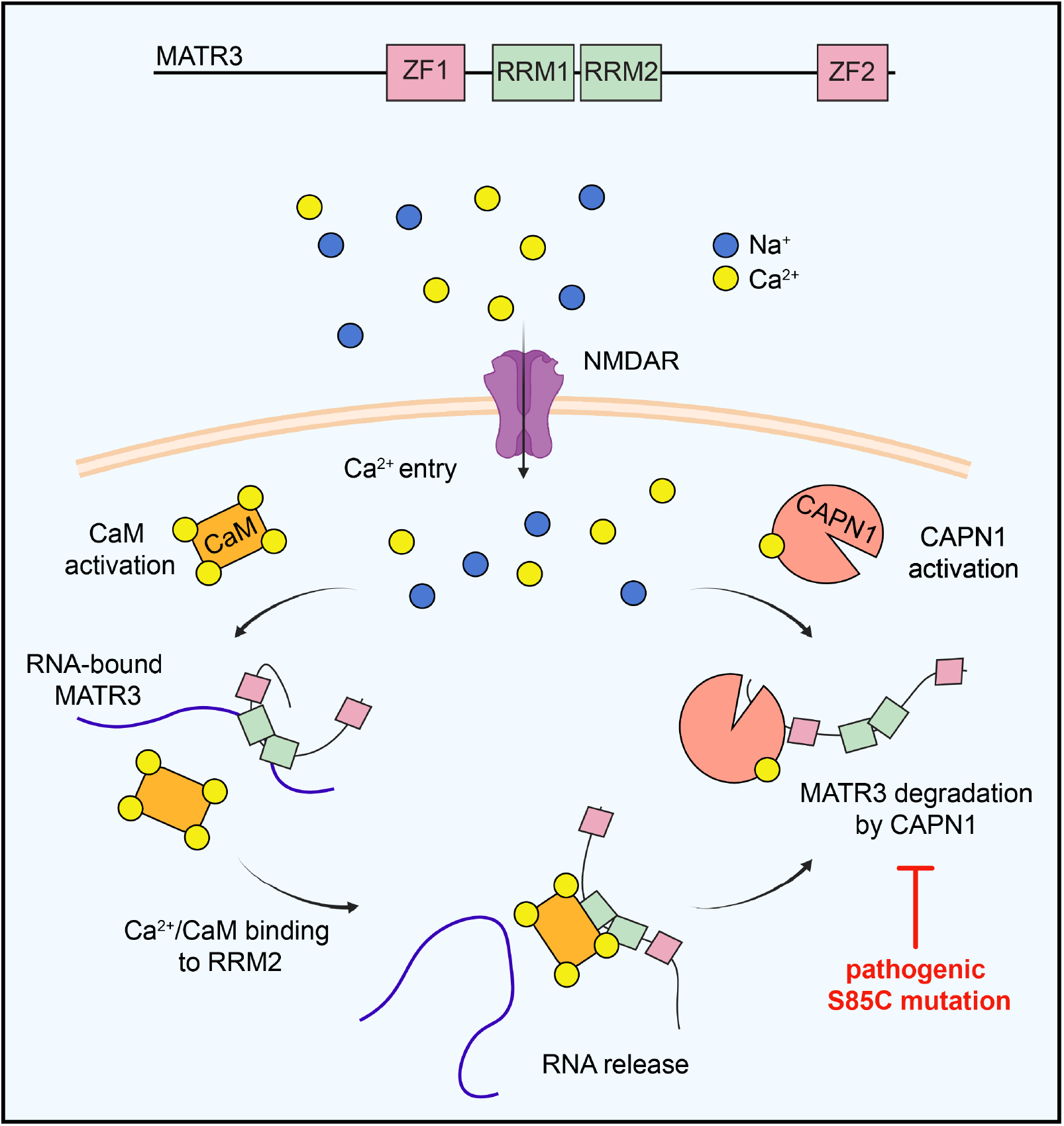
A model depicting regulation of MATR3 abundance and RNA binding in a Ca^2+^/CaM-dependent manner. MATR3 possesses two zinc finger (ZF) domains and two RNA recognition motifs (RRMs), with RRM2 being the dominant RNA-binding domain. Ca^2+^ influx through NMDARs activates calmodulin (CaM), which displaces bound RNA from MATR3 via overlapping interactions with RRM2. Simultaneously, Ca^2+^ activates calpain-1 (CAPN1), which is capable of degrading RNA-deficient MATR3. By virtue of its location within the predicted CAPN1 cleavage site, the pathogenic S85C mutation interferes with CAPN1-mediated MATR3 degradation.

Our results in cortical neurons conflict with those observed in cerebellar granule neurons, in which PKA inhibition by H89 completely blocks MATR3 degradation in response to NMDA and prevents MATR3 phosphorylation detected with a PKA phosphosubstrate antibody^36^. Surprisingly, H89 had little effect on MATR3 phosphorylation, implying the presence of an alternative MATR3 kinase acting on the consensus PKA motif (RRXpS/T) recognized by the PKA phospho-substrate antibody^62,63^. The identity of this kinase that phosphorylates MATR3 after NMDAR activation remains unknown, as does the functional significance of this modification. The relatively subtle effect of H89 on MATR3 turnover in our experiments may also be due to additional PKA targets that affect Ca^2+^ dynamics and, indirectly, MATR3 clearance. PKA is itself activated by Ca^2+^/CaM through adenylyl cyclase activation^64–66^ and phosphorylates key residues on the intracellular portions of NMDARs, thereby making them more conductive and enhancing Ca^2+^ influx through a feedforward-type mechanism^67^,^68^.The blunted MATR3 clearance we observe upon PKA (Supplemental Fig. 1C-D) and CaM inhibition (Supplemental Fig. 4A-B) may arise not from a direct effect of these molecules on MATR3, but rather indirectly, by influencing the complex crosstalk between Ca^2+^ and Ca^2+^-dependent signaling pathways in neurons.

Extending our findings beyond primary neurons into an *ex vivo* system, we found that cerebellar Purkinje neurons—a cell type previously reported to display altered MATR3 metabolism as baseline and in disease models—likewise clear MATR3 in response to glutamatergic stimulation. These observations suggest that differences in Purkinje cell activity may partially explain disparities in MATR3 abundance noted in previous studies^43^. Future experiments may examine the behavioral threshold required to drive MATR3 clearance in Purkinje neurons, perhaps through motor tasks^69^. Intriguingly, a knock-in mouse homozygous for the S85C mutation exhibited markedly reduced MATR3 staining in Purkinje cells from an early timepoint followed by age-dependent Purkinje cell loss^44^. The mechanism for how this mutation affects MATR3 metabolism in Purkinje neurons remains unclear, however, as are its implications for human genetics since no *MATR3* mutation identified to date results in cerebellar degeneration or Purkinje cell loss.

We found instead that the rapid decrease over a matter of hours in MATR3 abundance after NMDA treatment is dependent on Ca^2+^, and furthermore, that elevated Ca^2+^ is sufficient to drive MATR3 degradation not only in NMDAergically mature DIV14-16 (Fig. 3.2F-G) but also in immature DIV4-5 (Fig. 3.2H-I) neurons, indicating a key role for Ca^2+^ downstream of NMDAR recruitment. Although increases in intracellular Ca^2+^ can activate several proteolytic pathways, including autophagy^70,71^ and the UPS^72,73^, we found that neither of these processes were responsible for MATR3 clearance. Rather, calpain inhibition fully blocked NMDA-triggered degradation, suggesting that calpains are the final effectors of MATR3 clearance.

A growing body of evidence implicates calpain cleavage of RBPs in the pathogenesis of ALS/FTD and related disorders. Activated calpains degrade survival motor neuron (SMN)^74^ and are capable of cleaving MATR3, FUS, and TDP43 *in vitro* and *in vivo*^48,75^. Calpain-mediated cleavage of TDP43 can be modulated through phosphorylation^76^, but whether posttranslational modifications of MATR3^77^ likewise tune its susceptibility to clearance are unknown. Overexpression of calpastatin, an endogenous calpain inhibitor, prevented cleavage of calpain targets while increasing motor function and survival in a transgenic mutant SOD1 mouse model of ALS^78^. Though the connection with human disease is still unclear, these observations highlight the importance of calpain-mediated protein turnover mechanisms for ALS pathogenesis and perhaps therapeutic purposes.

Ca^2+^ and calpains regulate RBPs beyond simply degrading them, though these mechanisms are less defined. Increases in cellular Ca^2+^ result in the cytoplasmic redistribution of TDP43 and the ALS-linked RBP FUS^79^, while this same stimulus drives the nuclear enrichment of CPEB4^80^, indicating that Ca^2+^ can bidirectionally regulate localization in a unique and selective manner. An RNAi screen of calcium signaling proteins in *Drosophila* demonstrated that calpain A is critical for TDP43 cytoplasmic localization after Ca^2+^ increase^81^, and degradation of nuclear pore complex components by Ca^2+^-activated calpains resulted in aberrant nucleocytoplasmic transport in mouse CNS^82^. These results suggest that calpains may modulate RBP localization as well as abundance. Even so, we failed to detect any changes in MATR3 distribution after NMDA treatment (Fig. 1C), suggesting that Ca^2+^- and calpain-mediated MATR3 regulation occurs primarily within the nucleus, without a change in MATR3 localization.

In testing MATR3 susceptibility to the two main calpains present in the CNS, we found that WT protein—but not the pathogenic S85C mutant—is a substrate of CAPN1 (Fig. 4.D-E). The S85C mutation is the most common disease-linked variant in *MATR3*, with a phenotypic spectrum that encompasses both ALS and distal myopathy with vocal cord and pharyngeal weakness (VCPDM)^14,83–85^. In patient tissue^13^ as well as cellular^61^ and animal^44^,^86^,^87^ models, this mutation reduces MATR3 solubility through unknown mechanisms. Resistance to CAPN1 may promote aggregation and insolubility of MATR3(S85C); alternatively, insolubility may itself interfere with CAPN1-mediated degradation. Like neurons, skeletal muscle cells are excitable and rely not only on Ca^2+^ levels for their function but also on CAPN1 activation as a proteolytic mechanism to control myofiber size and number^88^. Therefore, our findings may offer insight into the unique disease spectrum of the S85C mutation that encompasses motor neuron and skeletal muscle pathology.

Seeking to investigate rapid, non-degradative regulation after neuronal activity, we pursued the binding between Ca^2+^/CaM and a sequence on MATR3 overlapping its RRM2, experimentally confirming this for MATR3 and TDP43 but also finding that according to prediction algorithms many disease-associated RBPs have CaM-binding motifs in their RRMs. Functionally, we replicated the observation that Ca^2+^/CaM enriches in the nucleus after activation and found that this is associated with impaired MATR3 binding to its RNA substrates. A similar phenomenon of inhibited nucleic acid recognition through activated CaM binding has been reported for CPSF30 in *Arabidopsis^89^*, as well as the CNS-specific mammalian PCP4^90^. Our preliminary results for MATR3 and TDP43 combined with predictions for other RBPs suggest that this may be a widely applicable mechanism for tuning RBP function in response to neuronal activity.

In the work presented here, we have shown that glutamatergic stimulation and the resulting increase in Ca^2+^ results in suppression of MATR3 function through two complementary mechanisms. Within minutes of Ca^2+^ influx, Ca^2+^/CaM binding to MATR3 rapidly impairs its ability to bind RNAs. Over extended periods of time, activation of calpains results in MATR3 cleavage. What consequences these regulatory processes have for MATR3 gene targets, particularly those that encode activity-related proteins, are currently unclear, as are the implications of this Ca^2+^-mediated control for neuronal physiology and for RBP-linked neuromuscular disease. Nevertheless, we expect that continued investigations will offer insights into the basic biology of activity-dependent RBP regulation, which can in turn be leveraged for therapies targeting neurological disorders.

## Materials and methods

### Plasmids and expression vector delivery

For lentiviral expression plasmids, the FLAG-MATR3 open reading frame (ORF) was PCR amplified from pGW1 FLAG-MATR3-Dendra2 using primers that attached HpaI sites to the 5’ and 3’ end of the ORF. The resulting amplicon was inserted into the HpaI site of pLV-EF1a. CAPN1 (#60941) and CAPN2 (#60942) overexpression constructs were purchased from Addgene. Expression plasmids for pGW1 FLAG-MATR3(WT) and FLAG-MATR3(S85C) were generated by amplifying MATR3(WT) and MATR3(S85C) from pGW1 FLAG-MATR3-EGFP constructs^61^ using Q5 Hot Start High-Fidelity DNA Polymerase (New England Biolabs) and flanking primers that eliminated linker sequences and EGFP. A similar strategy was used for creating MATR3(ΔRRM2)-Dendra2.

HEK293T cells (ATCC CRL-3216) were transfected in a 6-well plate with 3μg/well of DNA with Lipofectamine 2000 (ThermoFisher) in accordance with the manufacturer’s instructions. For lentivirus experiments, viral particles were generated by the University of Michigan Vector Core and neurons transduced at DIV5-6 overnight. The next day, media was removed from cells, after which they were washed once in PBS before being placed in virus-free condition media until experimentation on DIV14-16.

### Primary cortical neuron culture and pharmacological treatments

Primary cortical neurons were isolated from E19-20 rats and plated as previously described^91^. Neurons were treated at DIV14-16 with 100μM NMDA (Sigma) or L-glutamate (Sigma) for 3h or at the doses and times detailed in Fig. 1. For blocking experiments, cells were cotreated with 100μM D-AP5 (Tocris), 10μM MG132 (Sigma), 20nM bafilomycin (Sigma), 100nM bortezomib (Sigma), or 20μM MDL28170 (ThermoFisher). For Ca^2+^ chelation and PKA inhibition experiments, neurons were pretreated for 30m with 2mM BAPTA (Cayman Chemical) or 20μM H89 (Tocris), respectively, prior to NMDA application. To increase intracellular Ca^2+^, ionomycin (ThermoFisher) was applied at 10μg/mL for 3h. Cerebellar slices were incubated with 500μM glutamate or vehicle for 3h prior to immunostaining. All pharmacological treatments were performed at 37°C.

### Immunoblot

Cells were washed in PBS and lysed in RIPA buffer (Pierce) supplemented with cOmplete protease inhibitor cocktail (Roche). For experiments investigating MATR3 phosphorylation, the lysis buffer was also supplemented with PhosSTOP phosphatase inhibitor (Roche). Resuspended cells were sonicated using a Fisherbrand Model 505 Sonic Dismembrenator (ThermoFisher) at 80% amplitude, 5s on/5s off for 2m. Lysates were cleared by centrifugation at 21,000x g at 4°C for 15m. 5-20μg protein were loaded onto 4-20% SDS-PAGE gels (Biorad) and run at 100V, after which proteins were transferred onto 0.2μm PVDF membranes (Biorad) at 100V for 2h at 4°C. Membranes were incubated in blocking buffer (3% BSA, 0.2% Tween-20 in Tris-buffered saline (TBST)) and then blotted with the following primary antibodies diluted in blocking buffer: rabbit anti-MATR3 N-terminus [1:1000, Abcam EPR10634(B)] for all blots except for Supplemental Fig. 2 as noted; mouse anti-GAPDH (1:1000, MiliporeSigma MAB374); rabbit anti-MATR3 C-terminus (1:1000, Abcam EPR10635(B)) for Supplemental Fig. 2B; mouse anti-ubiquitin (1:100, Santa Cruz sc-8017); rabbit anti-TDP43 (1:5000, Proteintech 10782-2-AP); rabbit anti-phospho-PKA substrate (1:1000; Cell Signaling #9624). The following day they were washed 3 x 5m in TBST and incubated with donkey anti-mouse 800 (1:10,000, LI-COR 925-32213) and donkey anti-rabbit 680 (1:10,000 LICOR 926-68073) in blocking buffer, for 1h at RT. Membranes were then washed again 3 x 5m in TBST and imaged on an Odyssey CLx Imaging System (LI-COR).

### Immunocytochemistry

Primary cortical neurons were fixed in 4% PFA in PBS supplemented with 2mM CaCl2 at RT for 10m, permeabilized with 0.1% Triton X-100 in PBS, and treated with 10mM glycine in PBS. They were then placed in blocking solution (0.1% Triton X-100, 2% fetal calf serum, 3% BSA in PBS) for 1h at RT, after which they were probed overnight at 4°C with the following antibodies diluted in blocking solution: rabbit anti-MATR3 [1:1000, Abcam EPR10635(B)]; chicken anti-MAP2 (1:1000, Novus Biologicals NB300-213); or mouse anti-calmodulin (1:1000, ThermoFisher MA3-917). The next day, samples were washed 3 x 5m in PBS and then incubated 1h at RT with the following secondary antibodies all diluted 1:1000 in blocking solution: donkey anti-rabbit Alexa Fluor 647 (ThermoFisher A-31573); goat anti-mouse Alexa Fluor 568 (ThermoFisher A-11031); and goat anti-chicken Alexa Fluor 488 (ThermoFisher A-11039). After 3 x 5m PBS washes, neurons were imaged using an Eclipse Ti inverted microscope (Nikon) with PerfectFocus, Semrock filters, Lambda 421 lamp (Sutter Instruments), and an Andor Zyla 4.2(+) sCMOS camera (Oxford Instruments), with custom BeanShell scripts controlling image acquisition and stage movements via μManager^92^. For confocal microscopy, neurons were mounted in ProLong Gold Antifade Mountant with DAPI (ThermoFisher P36935), and imaged with a Nikon A1 inverted point-scanning confocal microscope (Nikon) controlled by NIS-Elements software. For CaM N/C ratio determination, custom scripts were used to create nuclear and cytoplasmic ROIs using DAPI and MAP2 signal, respectively.

### Cerebellar slice culture and immunohistochemistry

Wild-type C57BL/6J mice were anesthetized by isoflurane inhalation. Slices were prepared using a VT1200 vibratome (Leica) at a thickness of 300μM as before^93,94^ and placed in prewarmed aCSF with 5% CO_2_ and 95% O_2_ for 30m before transferring to Neurobasal Medium (ThermoFisher). They were treated with vehicle or 500μM glutamate for 3h at 37°C, after which they were processed as before^95^. Briefly, slices were fixed in 4% PBS at 4°C overnight. The following day samples were permeabilized with 1% Triton X-100 in PBS for 1.5h, treated with 20mM glycine in PBS for 1h, blocked in 5% normal goat serum in PBS, and probed overnight at 4°C with rabbit anti-MATR3 [1:1000, Abcam EPR10635(B)] and mouse anti-calbindin (1:1000, MiliporeSigma C9848). After washing 3 x 5m in PBS, slices were incubated overnight at 4°C with goat anti-mouse Alexa Fluor 568 and goat anti-rabbit Alexa Fluor 488 (ThermoFisher A-11008), both diluted 1:500 in blocking solution. Stained slices were washed 3 x 5m in PBS and then mounted in ProLong Gold Antifade Mountant with DAPI. Slices were imaged using a 40X objective Nikon A1 inverted point-scanning confocal microscope, as above.

### Immunoprecipitation

DIV14-16 primary neurons treated with vehicle (DMSO), 100μM NMDA, 10μM MG132, or both compounds, then collected in PBS, lysed in RIPA buffer supplemented with protease and phosphatase inhibitors and 2mM of the deubiquitinating enzyme inhibitor N-ethylmaleimide (MiliporeSigma), and sonicated at 80% amplitude, 5s on/5s off for 2m in a Fisherbrand Model 505 Sonic Dismembrenator. Lysates were cleared by centrifugation at 21,000x g at 4°C for 15m, and resuspended in dilution buffer (50mM Tris, 10mM EDTA, 1.5M NaCl, 1% Na-deoxycholate, 10% Triton X-100, pH 8.0) before incubation overnight at 4°C with 1μg rabbit anti-MATR3 antibody per sample (ThermoFisher A300-591A) conjugated to Dynabeads Protein G (ThermoFisher). The following day beads were washed 3 x 5m in wash buffer (20mM Tris, 1% Triton X-100, 2mM EDTA, 150mM NaCl, 0.1% SDS, pH 8.1) before elution by heating in SDS-PAGE sample buffer at 95°C for 10m.

### CaM-sepharose pulldown

HEK293T cells were collected in PBS, resuspended in buffer (10mM Tris, 5mM MgCl2, pH 7.4) supplemented with cOmplete EDTA-free protease inhibitor (Roche), and lysed by passing through an 18G x 1 ½” needle and sonication at 80% amplitude, 5s on/5s off for 2m in a Fisherbrand Model 505 Sonic Dismembrenator. Cleared lysates were rotated overnight at 4°C with 100μL per sample of Calmodulin Sepharose 4B resin (Cytiva Life Science) in CaM binding buffer (50mM Tris HCl, 150mM NaCl, 2mM CaCl2, 1μM DTT, pH 7.5). The next day, beads were washed 3 x 5m in CaM binding buffer and eluted in CaM elution buffer with EGTA (50mM Tris HCl, 150mM NaCl, 2mM EGTA, 1μM DTT, pH 7.5) or BAPTA (50mM Tris HCl, 150mM NaCl, 2mM BAPTA, 1μM DTT, pH 7.5). For inactivation of CaM, CaM binding buffer was supplemented with 100μM W-7 (Tocris).

### UV-CLIP and RT-PCR

DIV14-16 primary neurons were pretreated with vehicle (DMSO) or 100μM W-7 for 30m prior to application of vehicle or 100μM NMDA for 3-5m before crosslinking at 254nm in a Stratalinker 2400 (Stratagene) at 1500mJ/cm^2^. The cells were then lysed in RIPA buffer supplemented with cOmplete protease inhibitor (Roche) and RNaisin ribonuclease inhibitor (Promega) and sonicated at 80% amplitude, 5s on/5s off for 2m using a Fisherbrand Model 505 Sonic Dismembrenator. Cleared lysates were incubated overnight at 4°C with 2μg rabbit anti-MATR3 antibody/sample (ThermoFisher A300-591A) conjugated to Dynabeads Protein G (ThermoFisher). The following day, beads were washed 1x in NaCl wash buffer (20mM Tris, 1% Triton X-100, 2mM EDTA, 150mM NaCl, 0.1% SDS, pH 8.1), 1x in LiCl wash buffer (10mM Tris, 1% NP-40, 1mM EDTA, 0.25% Na-deoxycholate, 250mM LiCl, pH 8.0), and 2x in TE buffer (10mM Tris, 1mM EDTA, pH 8.0).

Samples were then split for either protein analysis, for which MATR3 complexes were removed from beads by heating at 95°C for 10m in SDS-PAGE sample buffer, or for RNA analysis. For the latter, beads were resuspended in elution buffer (50mM Tris, 1mM EDTA, 150mM NaCl, 1% SDS, 50mM NaHCO_3_, pH 8.1) supplemented with RNasin (Promega) and incubated with proteinase K (New England Biolabs) for 2h at 50°C. RNA was extracted with TRIzol reagent (ThermoFisher) and chloroform, the suspension centrifuged at 21,000x g at 4°C for 15m, the aqueous phase removed, and RNA precipitated overnight at −20°C and resuspended in water by heating at 65°C for 10m. cDNA was generated using iScript Reverse Transcriptase Supermix (Bio-Rad) and qPCR was performed with 200nM each of forward and reverse primers and PowerUp SYBR Green Master Mix (ThermoFisher) according to manufacturer’s instructions.

### Statistical analysis

Statistical analyses were performed in Prism 7 (GraphPad) or R. Data were plotted using Prism 7, and significance determined via the two-tailed t-test. One-way ANOVA with Tukey’s post-test for more than two comparison or unpaired t-test for comparing two groups were used to assess for significant differences among protein abundances, RNA levels, nuclear/cytoplasmic ratios, and half-lives. Data are shown as mean ± SEM unless otherwise stated.

## Acknowledgements

This work was funded by NINDS 2R01-NS097542 (S.J.B.), NIA P30 AG072931 (S.J.B.), the family of Angela Dobson and Lyndon Welch, the Robert Packard Center for ALS Research (S.J.B.), NIGMS T32 GM007863 (A.M.M.), NINDS F31 NS110119 (A.M.M.), and NIGMS T32 GM007315 (J.W.W). Portions of Fig. 1, 2, 4, 5,and 6 were made with BioRender.

## Notes

### Competing Interest Statement

The authors have declared no competing interest.

