## Supplemental Figures and Methods for "Neuronal activity regulates Matrin 3 levels and function in a calcium-dependent manner through calpain cleavage and calmodulin binding"

### Supplemental figure 1

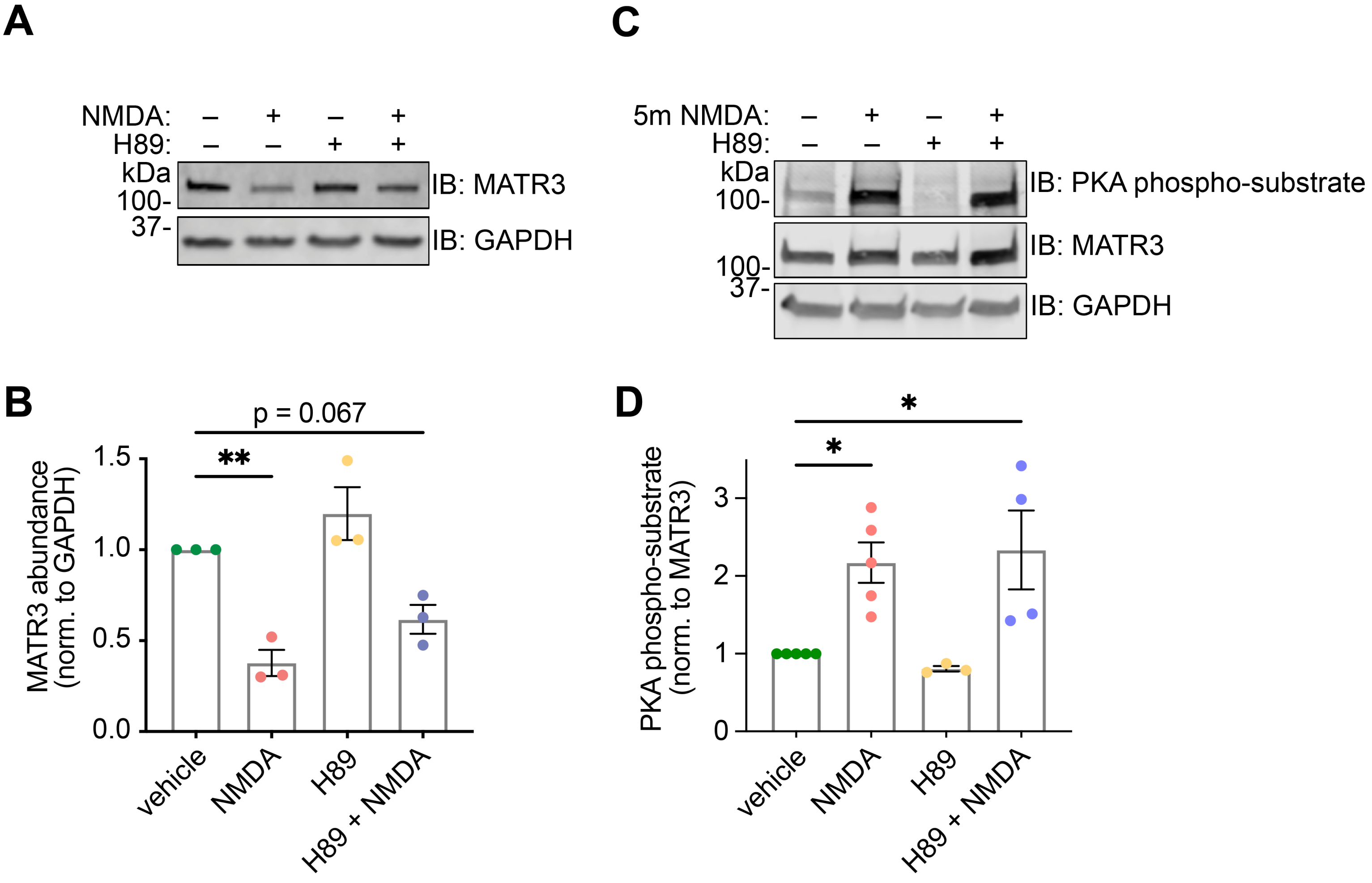

### Supplemental figure 2

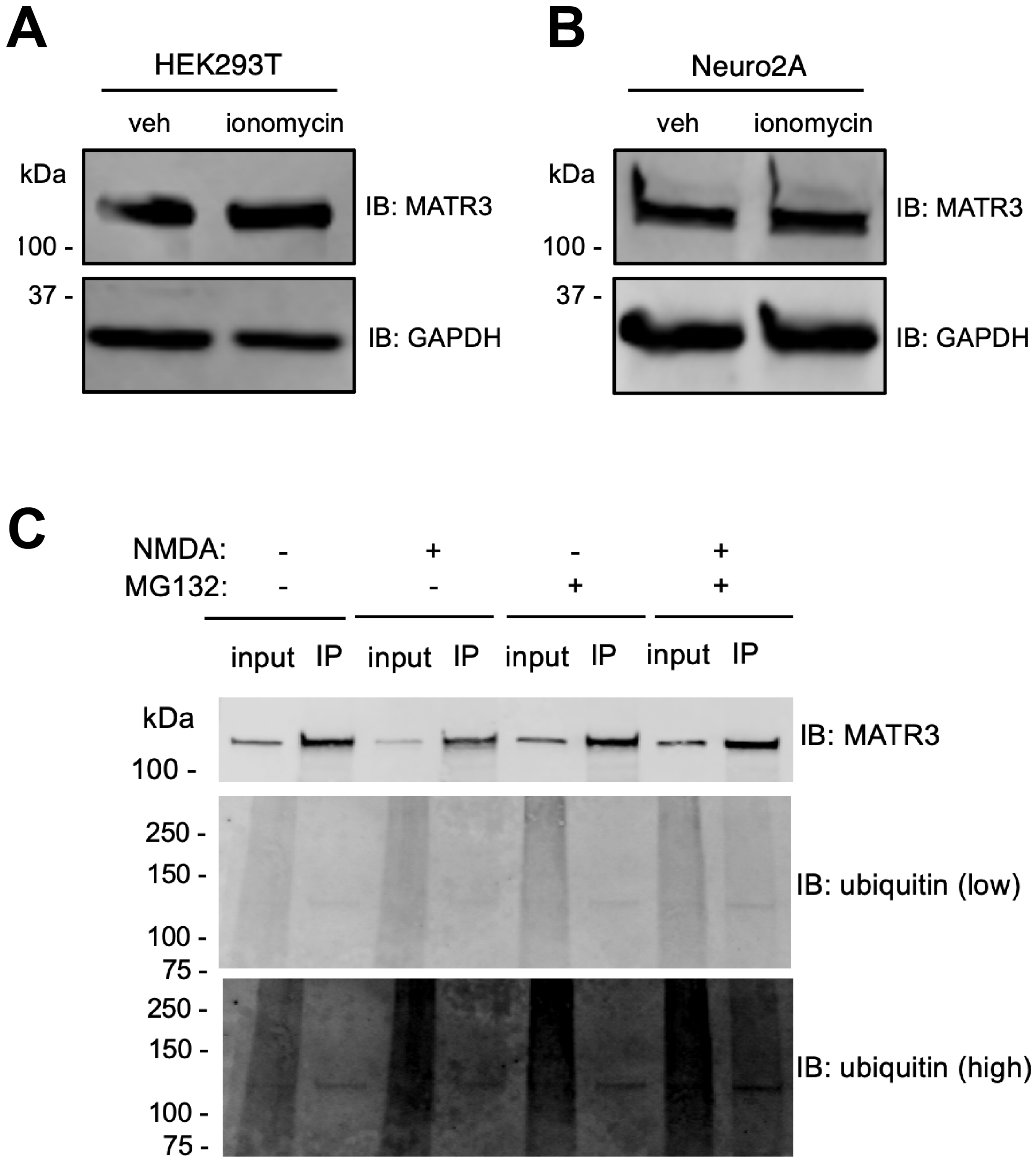

#### Supplemental figure 3

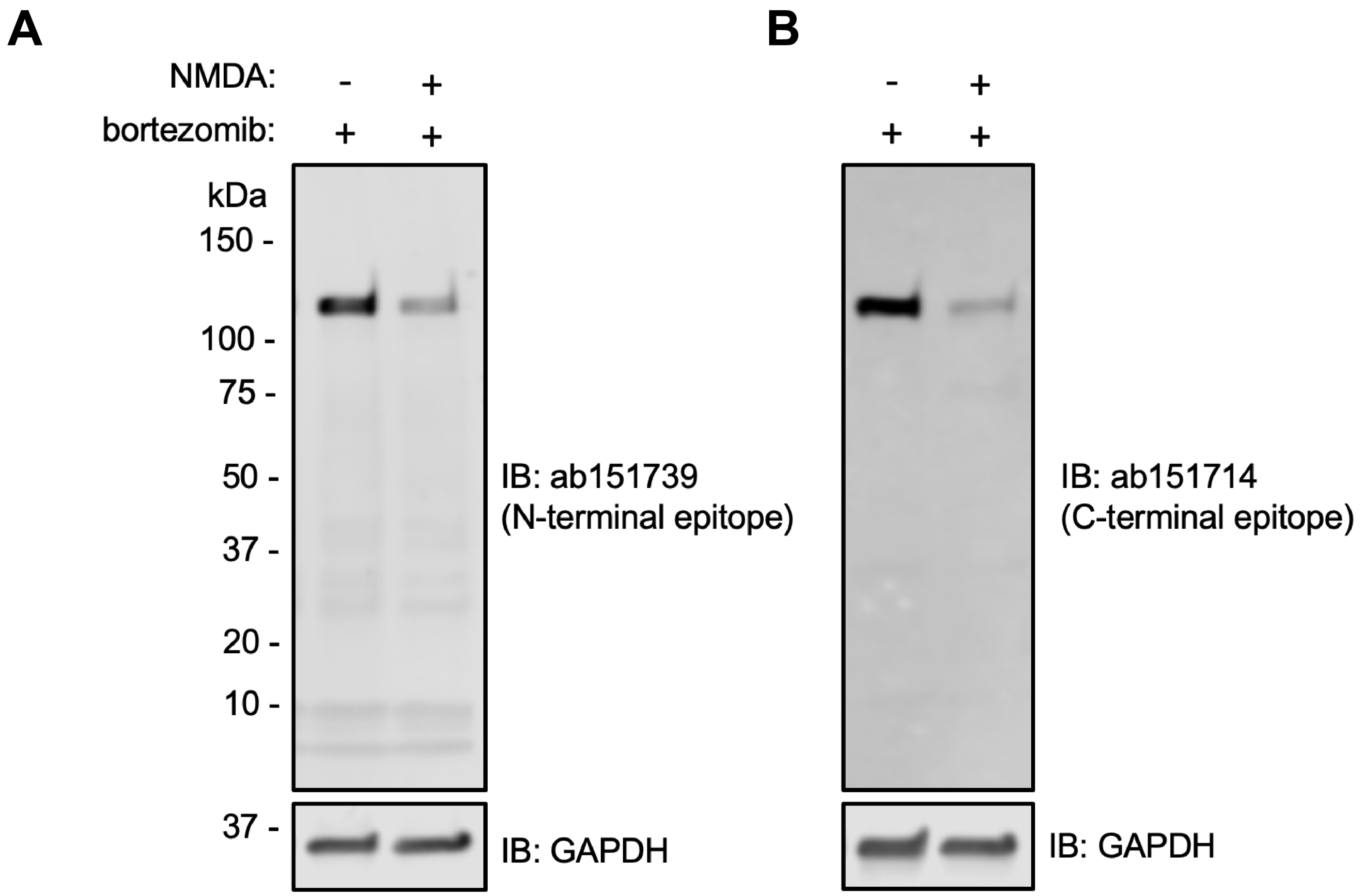

### Supplemental figure 4

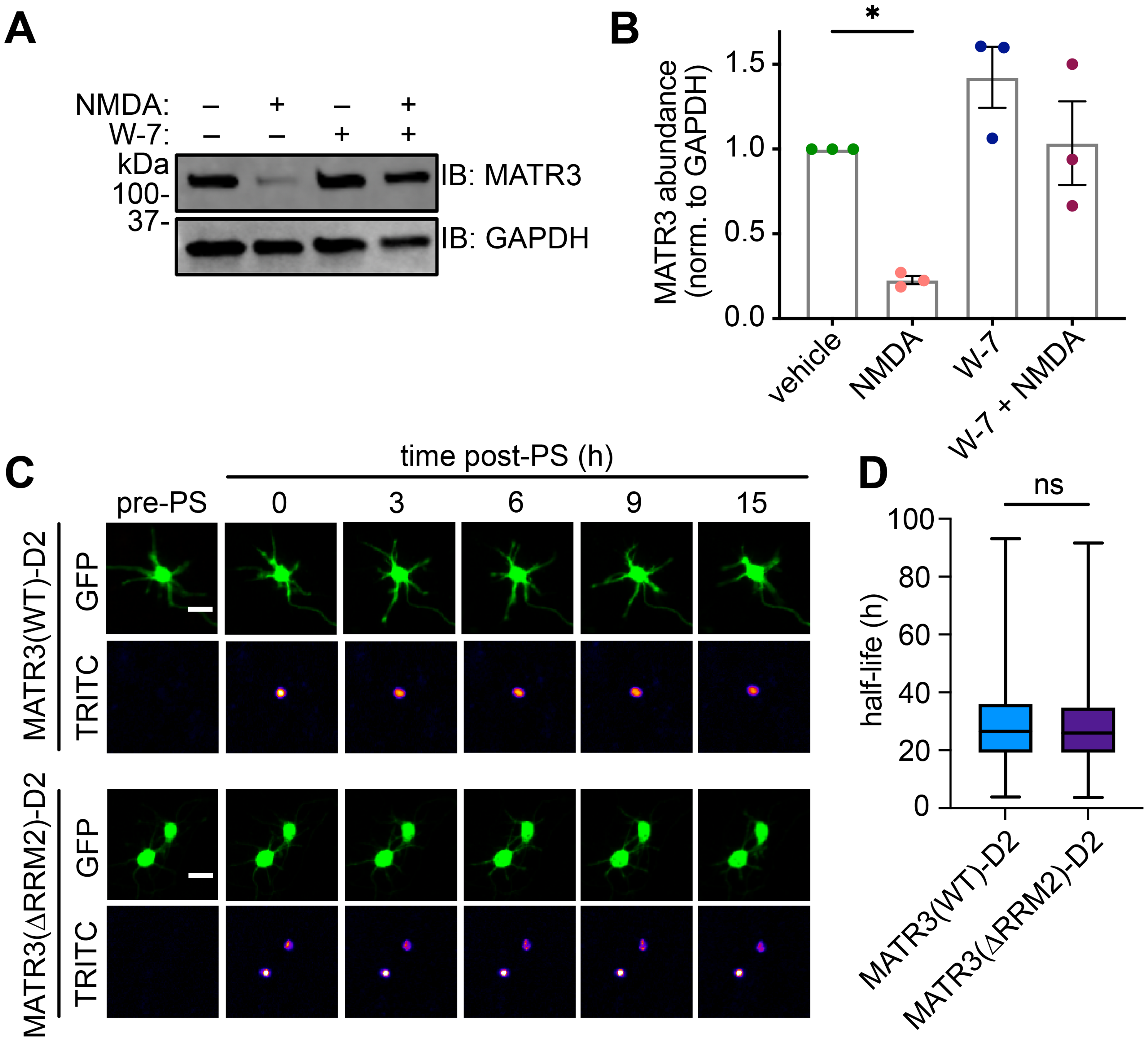

#### Supplemental methods

Optical pulse labelling and half-life determination

After imaging live cells with a Nikon Eclipse Ti inverted microscope as described above, neurons were identified in an automated manner based upon morphology, size, and fluorescence using customized Python scripts[1]. MATR3-Dendra2 was photoconverted with a 2s pulse of 405nm light, and single-cell protein half-lives were calculated by fitting log-transformed TRITC (RFP) intensity values measured from each cell at each time to a first-order exponential decay equation. Exponential curve fitting for determination of protein half-life was accomplished in R using customized scripts written for this purpose[1–4].

#### Supplemental figure legends

**Supplemental Figure 1. Inhibition of PKA blunts MATR3 degradation but does not block MATR3 phosphorylation.** (**A,B**) Treatment with the PKA inhibitor H89 trends toward partial inhibition of MATR3 degradation in response to NDMA treatment (compared to veh, n=3; NMDA, n=3, **p=0.0054; H89, n=3; p=0.45; H89 + NMDA, n=3; p=0.067). However, the increase in phospho-MATR3 5m after NMDA stimulation as detected with a general PKA phospho-substrate antibody is not blocked by H89, suggesting phosphorylation by another, H89-resistant kinase (compared to veh, n=5; NMDA, n=5, *p=0.033; H89, n=3, p=0.97; H89 + NMDA, n=4, *p=0.022).

**Supplemental Figure 2. NMDA-related MATR3 clearance occurs in a cell type-specific manner via a ubiquitin-independent pathway.** (**A,B**) Unlike in cortical neurons, MATR3 in HEK293T cells and Neuro2A cell lines is not degraded upon treatment with the ionophore ionomycin. (**C**) Proteasomal blockade with MG132 in the context of NMDA stimulation does not result in the accumulation of ubiquitinated, higher molecular weight MATR3 species, suggesting ubiquitin-independent means of MATR3 degradation.

**Supplemental Figure 3. Lower molecular weight MATR3 cleavage products are not detected in NMDA-treated neurons.** (**A,B**) Antibodies directed towards both the N- and C-termini of MATR3 cannot detect lower molecular weight MATR3 species even in the context of proteasomal blockade with bortezomib, suggesting that MATR3 fragments are highly unstable and are cleared through non-proteasomal mechanisms.

**Supplemental Figure 4. CaM inhibition blocks NMDA-triggered MATR3 degradation, but RNA binding-deficient MATR3 is not destabilized.** (**A,B**) Treatment with the CaM inhibitor W-7 blocks MATR3 reduction after NMDA treatment (compared to veh, n=3; NMDA, n=3, *p=0.030; W-7, n=3, p=0.28; W-7 + NMDA, n=3, p>0.99). (**C**) MATR3(WT) and MATR3(ΔRRM2) fused to the green-to-red photoconvertible protein Dendra2 were used in optical pulse labelling (OPL) experiments, with the decrease in red signal (TRITC) following photoswitching (PS) used to determine half-lives on an individual cell basis. (**D**) RRM2 deletion did not affect MATR3-Dendra2 half-life, arguing against destabilization of MATR3 variants that cannot bind RNA (MATR3(WT)-D2, n=495; MATR3(∆RRM2)-D2, n=701, p=0.47). Scale bars in (**C**), 20μm.

**Supplemental Table 1. Primer sequences for RT-PCR targets**

| Target | Primers | Sequences |
|---|---|---|
| *Matr3* | Forward | 5’-GCC ACC TCC TTC ATT TCA TCT-3’ |
| | Reverse | 5’-CTG GTT TCC ACT CTG CCT TT-3’ |
| *Gapdh* | Forward | 5’-AAG GTG AAG GTC GGA GTC AA-3’ |
| | Reverse | 5’-AAT GAA GGG GTC ATT GAT GG-3’ |
| *Tubb3* | Forward | 5’-GGC ATG GAT GAG ATG GAG TT-3’ |
| | Reverse | 5’-CTC CTC GTC GTC ATC TTC ATA C |
| *Lsamp* | Forward | 5’-CGG GAT GAC ACC AGG ATA AAC-3’ |
| | Reverse | 5’-CAC AGG TAT AGT TGC CGT AGT G-3 |
| *Grm7* | Forward | 5’-CAA GGA TCT GTG TGC TGA CTA C-3’ |
| | Reverse | 5’-GGT TCC AGC ACT ACC ATT GA-3’ |
| *Ftx* | Forward | 5’-GGC ACT TTG GGT CCC TAT ATC-3’ |
| | Reverse | 5’-CAG GTT TGT GCG TAT GTG TAA G-3’ |
